# Quantitative description of six fish species’ gut contents and abundances of their prey in the benthic food web of the Baltic Sea (1968-1978)

**DOI:** 10.1101/2023.12.08.570787

**Authors:** Benoit Gauzens, Gregor Kalinkat, Ana Carolina Antunes, Thomas Boy, Eoin J. O’Gorman, Ute Jacob, Malte Jochum, Susanne Kortsch, Benjamin Rosenbaum, Ludmilla Figueiredo, Ulrich Brose

## Abstract

The dataset presents a compilation of stomach contents from six demersal fish species from two functional groups inhabiting the Baltic Sea. It includes detailed information on prey identities, body masses, and biomasses recovered from both the fish’s digestive systems and their surrounding environment. Environmental parameters, such as salinity and temperature levels, have been integrated to enrich this dataset. The juxtaposition of information on prey found in stomachs and in the environment provides an opportunity to quantify trophic interactions across different environmental contexts and investigate how fish foraging behaviour adapts to changes in their environment, such as an increase in temperature. The compilation of body mass and taxonomic information for all species allows approaching these new questions using either a taxonomic (based on species identity) or functional trait (based on body mass) approach.

## Background and summary

In nature, the body mass of species is a fundamental trait related to the physiology of organisms^1^, the structure of communities, and their ability to persist over time^2,3^. Therefore, due to its capacity to explain a wide range of processes across different biological scales, it serves as a central element in numerous ecological studies and it is a key parameter for modelling, understanding and predicting the future of communities^4,5^. Specifically, the body mass relationship between predators and their prey (or predator-prey body mass ratio, PPMR) determines the occurrence and the strength of trophic interactions^6,7^, which has profound consequences for the stability of food webs and the coexistence of species within ecosystems^8^, as well as their response to environmental perturbations such as climate change^9^.

To explore these relationships, species-level food webs are often constructed using known feeding relationships, while body size information is independently added, typically based on species-level averages^10,11^. However, even within a predator-prey species pair, significant ontogenetic diet shifts can occur, leading to considerable variation in individual-level body size distributions. This important variability, obscured by the use of averaged values, is crucial because intra-specific variation in body mass can be greater than interspecific differences. Many methods for identifying individual-level trophic relationships, apart from time-intensive direct observation, cannot provide information on the size of individual prey items that are ingested (e.g. for terrestrial invertebrates^12^). Although there are very few existing datasets where predator and prey sizes were identified for invertebrates(e.g.^13^), more of such datasets exist for (commercially relevant) fish species^14–16^.

These datasets primarily focus on consumers and their stomach contents without providing information on the availability of their resources in the environment (but see^15^) and under different environmental conditions. This limitation hampers our ability to infer how consumers adapt their foraging behaviour to local environmental conditions and varying resource densities. It is a significant drawback of the existing databases, as the foraging behaviour has been shown to respond to biotic and abiotic constraints. Regarding abiotic conditions, species can adjust their diet depending on the underlying habitat complexity^17^ or temperature^18^. In terms of biotic factors, consumers can adapt their foraging strategy in response to the presence of their own predators^19^, to minimise competition with similar species^20^, and to adapt to the abundance of their own prey^21^.

Understanding the response of species foraging behaviour to local conditions is crucial for comprehending species coexistence and predicting community responses to climate change (Gauzens et al. in prep). For example, the adaptation of species foraging behaviour to the abundance of their prey is considered a solution to resolve May’s paradox^22^. Mathematically, large and complex communities are predicted to be unsustainable, which contradicts the general observation of species-rich natural ecosystems. Incorporating adaptive foraging behaviour into mathematical models has aligned theoretical predictions with empirical observations^23^. However, when aiming at predicting the coexistence of communities, most theoretical approaches rely on the assumption of optimal foraging theory: consumers will actively select their prey in order to maximise their net energetic income.

So far, we still lack a solid framework describing how consumer foraging behaviour—and, consequently, predator-prey body mass ratio (PPMR)—changes depending on environmental contexts. Our dataset will facilitate the analysis of these questions. It compiles the body mass of 6 different consumer fish species together with the identities, body masses and biomasses of the prey items found in their digestive systems and their environment. This data was sampled using systematic and standardised bottom trawls carried out year-round between 1968 and 1978 in Kiel Bay, located in the Baltic Sea (see Fig. 1). In total, the database comprises detailed dietary information based on stomach contents from 22,185 individuals of six commercially and ecologically important demersal fish species. These include two species of Gadiformes: “*Gadus morhua*” (Atlantic cod) and “*Merlangius merlangus*” (Whiting), as well as four species of Pleuronectiformes: “*Limanda limanda*” (common dab), “*Pleuronectes platessa*” (European plaice), “*Platichthys flesus*” (European flounder), and “*Hippoglossoides platessoides*” (American plaice).

**Figure 1:**
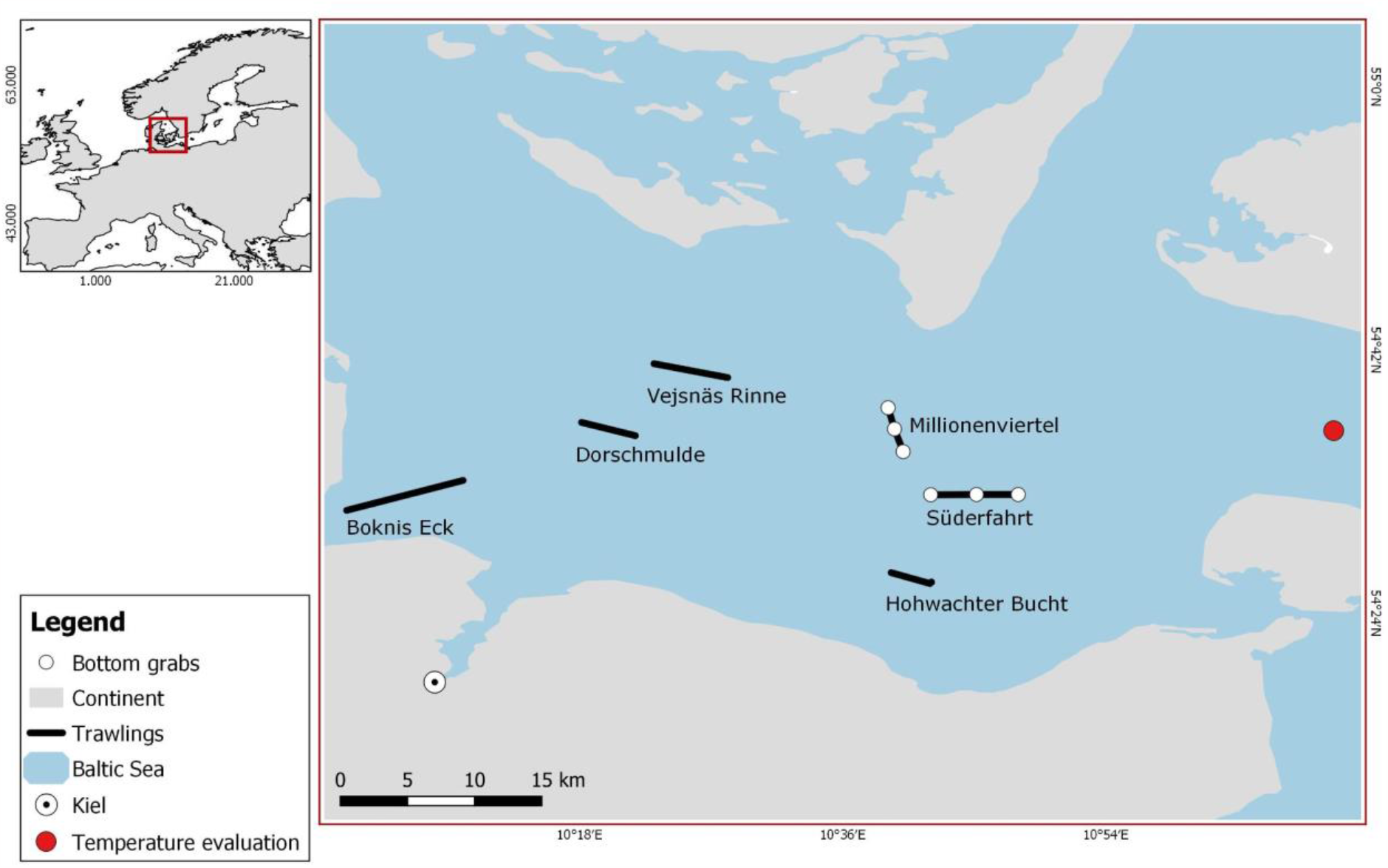
Location of bottom trawls (lines) and bottom grab stations (numbered dots) in Kiel Bay. The red point defines the station for temperature measurements (54° 35.894 N; 11° 08.895 E). Adapted from Arntz et al. 1974.

This dataset was enriched with information on local environmental conditions (local sea temperature, salinity). This joint availability of abiotic and biotic information enables an unprecedented level of understanding of how species foraging behaviour and individual PPMR vary depending on ecological contexts.

## Methods

Kiel Bay is located in the Western Baltic Sea, which is a marginal sea connected to the North Atlantic and considered the largest brackish sea in the world. It is a rather low-productivity ecosystem with low biodiversity due to its recent glacial history and its strong salinity gradients that only a few species can tolerate^24,25^. The core of the Kiel Bay database comprises detailed dietary information based on stomach contents from 5546 adult and sub-adult fish individuals of 6 species (Fig. 2), as well as the record of fish prey in the environment. The samplings of fish stomach contents and prey in the environment were conducted between the years 1968 and 1978 under the direction of Wolf Arntz. None of the authors participated in the data collection. A subset of this dataset was published in German language to describe the stomach contents of 2 of the fish species, “*Gadus morhua”*^29^ (Atlantic cod) and “*Limanda limanda*”^27^ (common dab).

**Figure 2:**
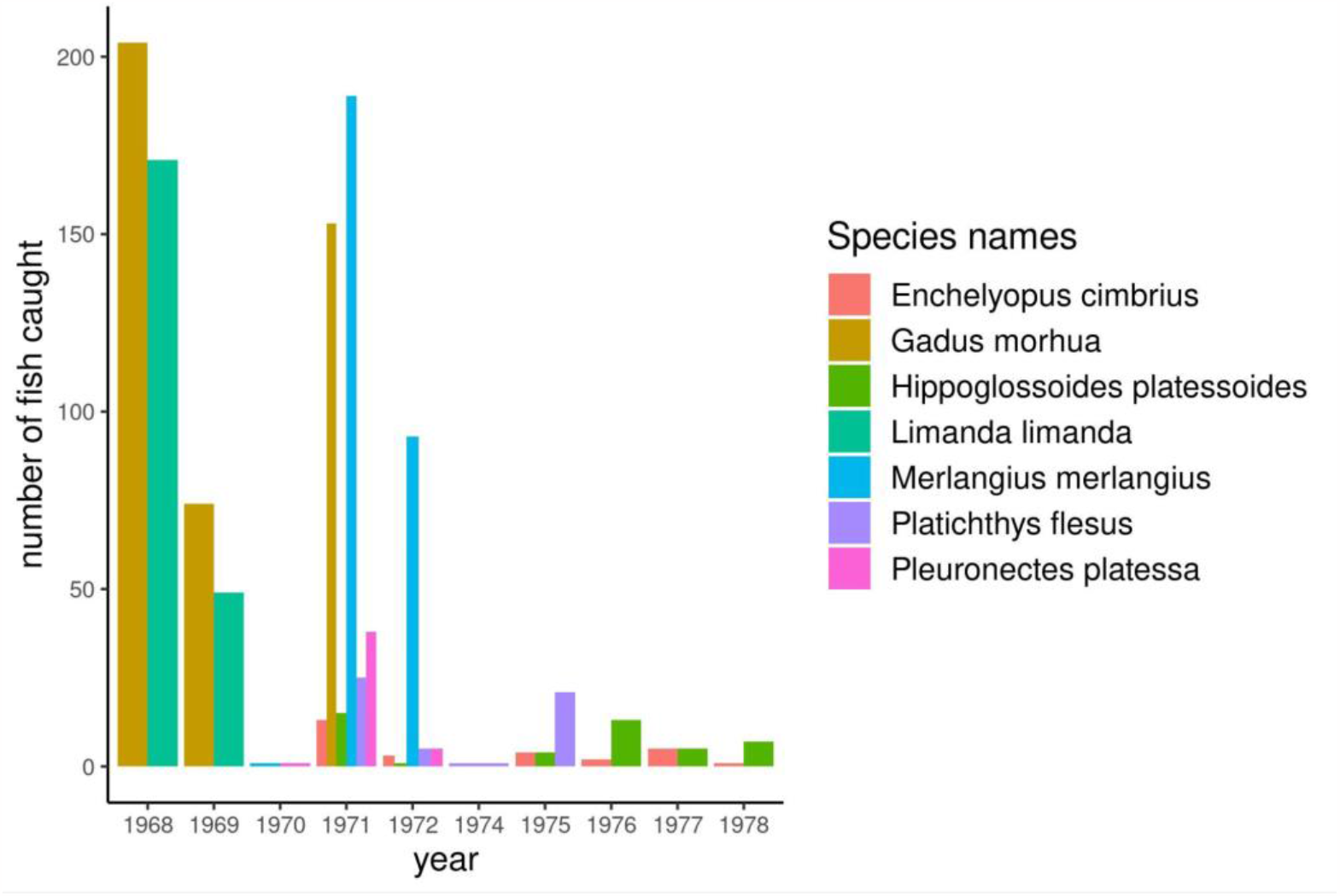
Distribution of fish analysed over the period covered by the database. Colours represent different fish species, and the y-axis shows the number of individuals harvested for each trawling.

We have compiled together the information on the 6 fish species as well as the environmental distribution of prey and digitized this historical dataset into an SQL database.

Fish species were classified into two functional groups based on their body shape and habitat use: (i) fusiform and benthopelagic species: “*Gadus morhua*” (Atlantic cod), “*Merlangius merlangus*” (Whiting); (ii) flat and demersal species (Pleuronectiformes): “*Limanda limanda*” (common dab), “*Pleuronectes platessa*” (European plaice), “*Platichthys flesus*” (European flounder), and “*Hippoglossoides platessoides*” (American plaice). This body-shape characteristic also corresponds to specific foraging behaviour^26^. Trait information for the fish species “Enchelyopus cimbrius” is also available but its stomach contents were not analysed.

The fish individuals were sampled using systematic bottom trawls carried out between 1968 and 1978 (Fig 3) and consisting of 24 hours of fishing with nets of 12 mm mesh size.

**Figure 3:**
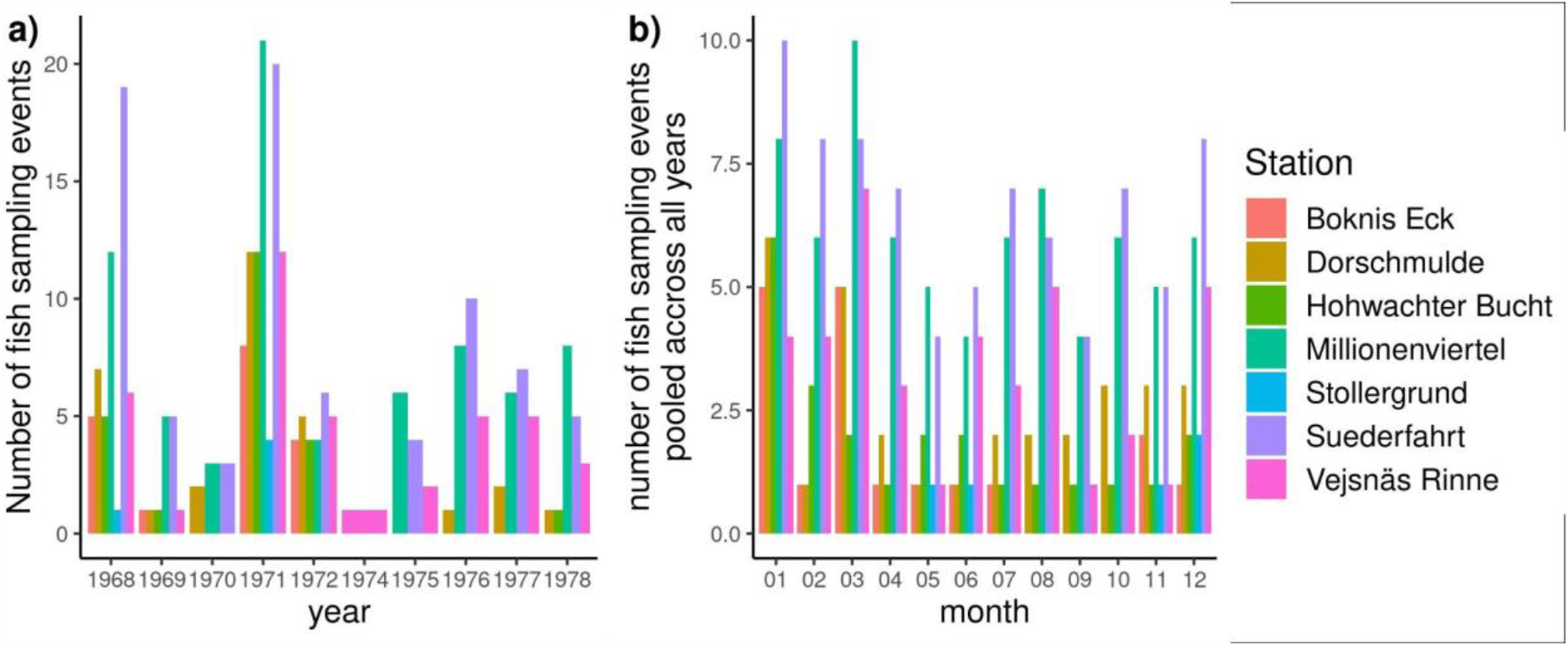
Number of fish sampling events. (a): per year, (b): per month, pooled across all years.

**Figure 3:**
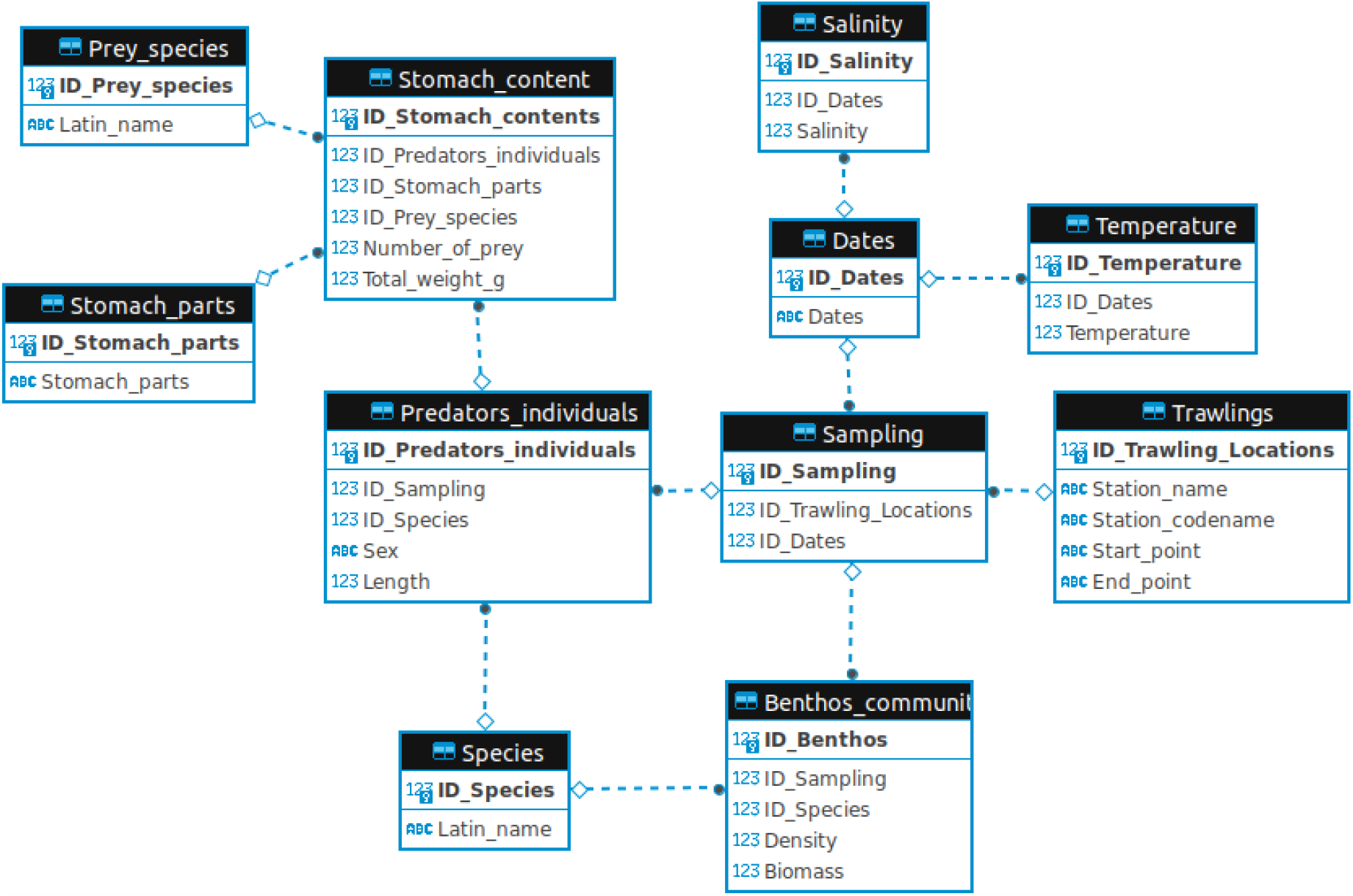
Entity relationship diagram representing the structure of the database. Variables in bold represent the primary keys (i.e. unique identifier for each datapoint) for each table.

According to the information available^27–30^, some substantial variation occurred in the size of the trawling net (15m or 24m width), and the number of hauls performed (between 8 and 12 per 24 hours). As such, it is impossible to ensure that the sampling intensity was fully standardised and any diversity analysis (either on taxonomic or trait diversity) should consider using rarefaction analyses. The body lengths of fish were measured and rounded to the nearest integer in cm. Stomach contents of all collected fish were identified to the highest taxonomic resolution possible (mostly species or genus level) and drained wet mass has been determined where possible. Fish lengths were measured (and rounded to the closest cm).

Samples from fish stomachs were fixed in 10% formalin and samples from benthos communities in 4% formalin. Both were weighed in the lab after the fishing trips. Note that today we know that these methods may potentially influence weight measurements. More specifically they can create inaccuracies because weight loss can vary for different organisms and depending on the time between preservation and weight determination (Wetzel et al 2005). Moreover, due to different digestion times for different groups of organisms this can potentially amplify already existing biases in the detectability of prey items from stomach contents. These shortcomings have to be taken into account for various kinds of analyses, particularly when comparing our data set with other data sets where samples were processed differently. Hence, the database includes body size data for all fish (i.e. predators), but also for prey items from the stomach contents. Items that were not identifiable were discarded; when body mass estimation was impossible it was coded as NA in the dataset. In addition, we were able to complement the stomach-content data with abundance and body mass information on the benthic invertebrate (i.e. prey) fauna from the surrounding environment to the database, which was sampled independently at the trawling locations using 0.1 m^2^ van Veen grabs^31^ (see^32,33^ for detailed procedure). Bottom trawls and bottom grabs (i.e. macrobenthos samplings with van Veen grabs) were always carried out in the same locations following a standardised protocol: 40 Kg Veen grabs with 0.5 mm mesh size were used. At each sampling date, 3 grabs were taken in each location. Samples were flushed onto a ground sieve with 0.9 mm mesh size to reduce the loss of small animals.

Molluscs were weighed with shells, and polychaetes without tubes. A detailed map of both trawling areas and bottom-grab locations is provided in Fig 1 (adapted from^34^). There, each trawling route within the given locations is delineated as a straight line of several kilometres (roughly, between 5 and 15 km). Bottom grab sampling locations were located directly along two of the trawling routes. In total, 6 sampling sites (3 for each of the two trawling routes) were used for the bottom grab sampling, each site being sampled several times during the period covered by our dataset (Fig. 4). Data from sampling sites were pooled together if they were harvested on the same date and corresponded to the same trawling route.

**Fig. 4:**
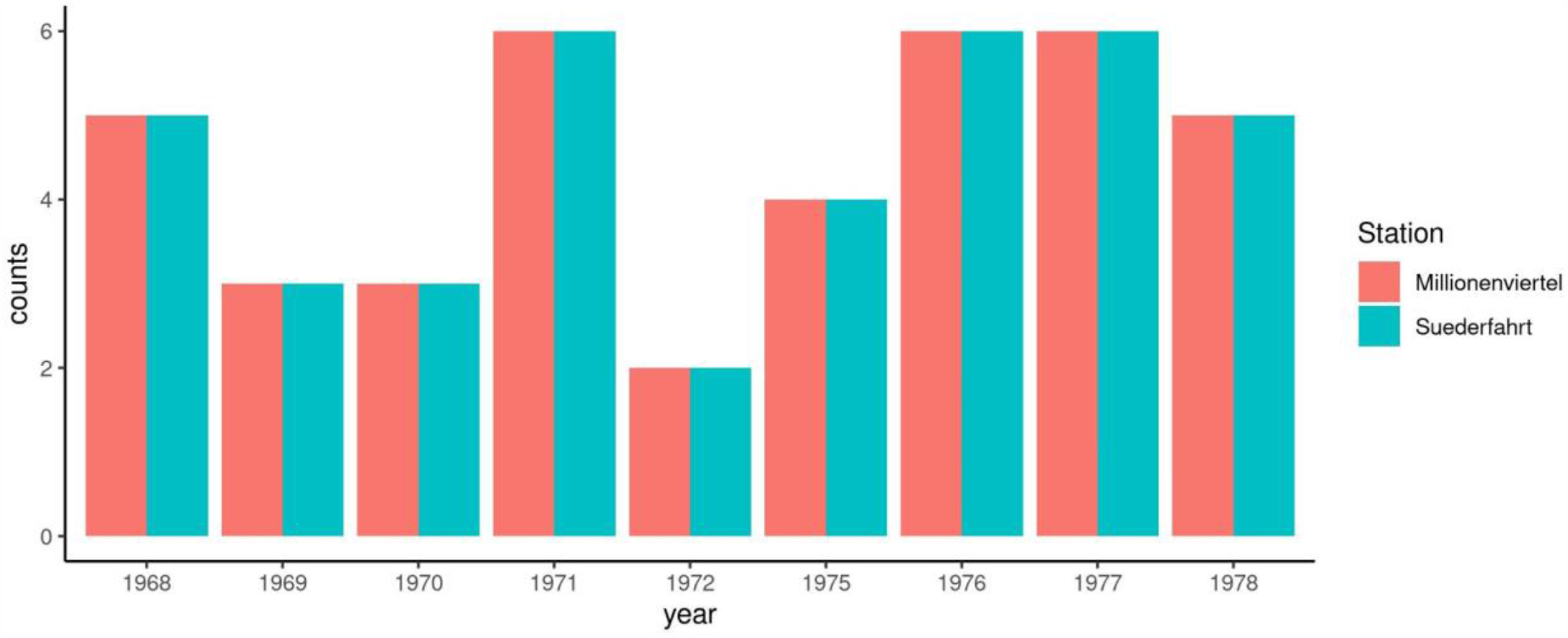
Number of benthos samplings performed per year for the two stations.

**Figure 4:**
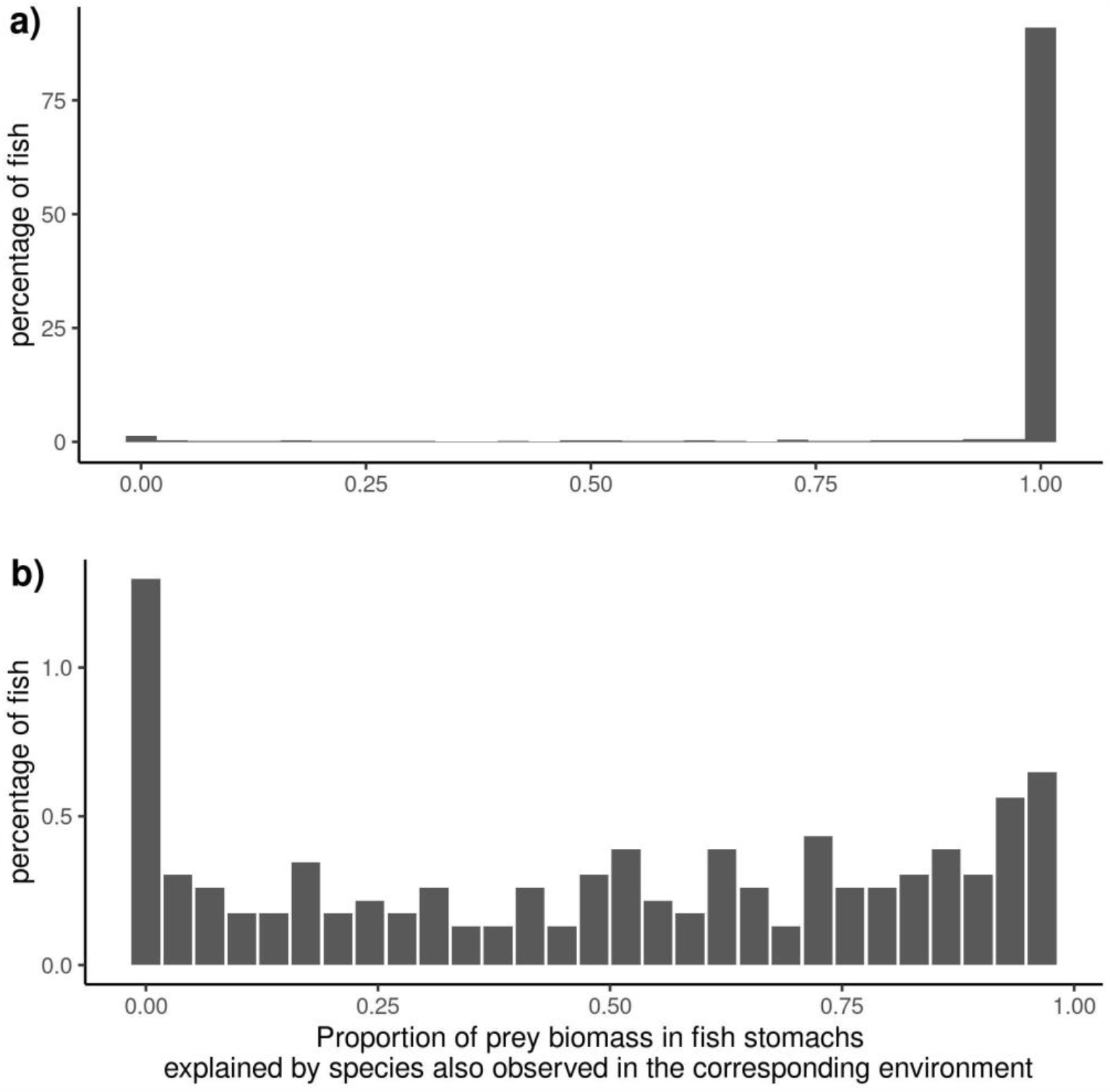
Correspondences between prey species observed in fish stomachs and in their corresponding environments in the full dataset (panel a) and focusing on fish for which less than 98% of prey biomass found in stomachs correspond to species also found in their environment (panel b). The x-axis represents the proportion of the prey biomass in fish stomachs that comes from species also found in the environment. A value of 1 means that all prey species found in fish stomachs were also found in the environment, and a value of 0.5 means that only 50% of the prey biomass in fish stomachs correspond to species also found in the environment

The temperature measurements represent monthly averages at 20 m depth in the Fehmarnbelt, position: 54°35,894N, 11°08,895E (Deutsches Hydrographisches Institut, DHI). Measured surface salinity data was extracted from^35^, where they are reported for the area between 54° N and 55° N, 10° E and 11° E.

### Data Records

The database is stored as an SQLite database on the iDiv biodiversity data portal, and is available at: <https://doi.org/10.25829/idiv.3547-rtgq13>. The different tables included in the database are related as depicted in the entity relationship diagram provided in Fig. 3. The database comprises 11 tables in total. The content of the different data tables, as well as variable types and units, can be found in Table 1):

**Table 1:** Formal descriptions of variables, their types and units for the fields of the different tables.

| ***Table*** | ***Field*** | ***Data***<br>***Type*** | ***Description*** |
| --- | --- | --- | --- |
| Prey_species | ID_Prey_species | INTEGER | PRIMARY KEY |
|  | Latin_name | TEXT | Binomial name (genus species) |
| Species | ID_Species | INTEGER | PRIMARY KEY |
|  | Latin_name | TEXT | Binomial name (genus species) |
| Dates | ID_Dates | INTEGER | PRIMARY KEY |
|  | Dates | DATE | date at the format YYYY-MM-DD |
| Trawlings | ID_Trawling_Locations | INTEGER | PRIMARY KEY |
|  | Station_name | TEXT | Local name of the station |
|  | Station_codename | TEXT | Code associated to the station |
|  | Start_point | TEXT | Geographic coordinates for the trawling starting point |
|  | End_point | TEXT | Geographic coordinates for the trawling end point |
| Sampling | ID_Sampling | INTEGER | PRIMARY KEY |

|  | ID_Trawling_Locations | INTEGER | FOREIGN KEY to the Trawlings table |
| --- | --- | --- | --- |
|  | ID_Dates | INTEGER | FOREIGN KEY to the Dates table |
| *Table* | *Field* | *Data Type* | *Description* |
| Benthos_community | ID_Benthos | INTEGER | PRIMARY KEY |
|  | ID_Sampling | INTEGER | FOREIGN KEY to the Sampling table |
|  | ID_Species | INTEGER | FOREIGN KEY to the Species table |
|  | Density | REAL | Density of the species, in individuals per m<sup>2</sup> |
|  | Biomass | REAL | Total biomass of the species in g.m<sup>-2</sup> at the ID_sampling event. |
| Predators_individuals | ID_Predators_individuals | INTEGER | PRIMARY KEY |
|  | ID_Sampling | INTEGER | FOREIGN KEY to the Sampling table |
|  | ID_Species | INTEGER | FOREIGN KEY to the Species table |
|  | Sex | TEXT | Sex of the individual |
|  | Length | REAL | Body length of the individual, in cm |
| Stomach_parts | ID_Stomach_parts | INTEGER | PRIMARY KEY |
|  | Stomach_parts | TEXT | Names of stomach parts used in the dataset |
| Stomach_content | ID_Stomach_contents | INTEGER | PRIMARY KEY |
|  | ID_Predators_individuals | INTEGER | FOREIGN KEY to the Predators_individuals table |
|  | ID_Stomach_parts | INTEGER | FOREIGN KEY to the Stomach_parts table |
|  | ID_Prey_species | INTEGER | FOREIGN KEY to the Prey_species table |
|  | Number_of_preys | INTEGER | Total number of individuals from the prey species |
|  | Total_weight_g | REAL | Total biomass (in g) of the corresponding prey species |

| *Table* | *Field* | *Data Type* | *Description* |
| --- | --- | --- | --- |
| Temperature | ID_Temperature | INTEGER | PRIMARY KEY |
|  | ID_Dates | INTEGER | FOREIGN KEY to the Dates table |
|  | Temperature | REAL | Temperature value in °C |
| Salinity | ID_Salinity | INTEGER | PRIMARY KEY |
|  | ID_Dates | INTEGER | FOREIGN KEY to the Dates table |
|  | Salinity | REAL | Salinity values, in Practical Salinity Units (PSU) |

### Species

contains the binomial names of species found in the environment (i.e. fish species and benthic species).

### Prey_species

contains the binomial names of species found in fish stomachs.

### Dates

a list of dates corresponding to events in the database (sampling, temperature, or salinity measurements).

### Temperatur

water temperature in Kiel Bay for a given date (referred to as ID_Dates).

### Salinity

water salinity in Kiel Bay for a given date (referred to as ID_Dates). Salinity was estimated at geographic coordinates 10°-11° East/54°-55° North. Extracted from^35^.

### Stomach_parts

the list of stomach parts in which the prey items were identified.

### Trawling

names and codes (local shortened names) of the locations where the sampling was done, as well as geographic coordinates of the starting and ending points. Bottom grabs for benthic communities were performed in the same locations.

### Sampling

a sampling occasion (can be either fish or benthic communities) is defined by a place (ID_Trawling_locations) and a date (ID_Dates).

### Benthos_community

the benthic community (i.e. a set of species, referred to as ID_Species) corresponding to a sampling event (ID_Sampling) and including information on densities and total biomass.

### Predator_individuals

individuals from a fish species (obtained from ID_species) captured at a given sampling event (ID_Sampling). Each individual has sex and body length (in cm) information.

### Stomach_content

the prey (ID_Prey_species) found in fish (ID_Predator_individuals) stomachs, from different stomach parts (ID_Stomach_parts). Contains information on the number of prey individuals of each prey species (Number_of_prey) as well as the total biomass of each prey species (Total_weight_g) for fish individual and stomach parts.

## Technical Validation

**Some prey items were not identifiable in fish stomachs are referred in the dataset as: “Siphons”, “Other Polychaeta”, “Other Polychaeta”, “Unidentified crustacean remains”, “Other Gastropoda”, “Various Groups” and “Fish offal”. These unidentified items represent 0.67% of all the recorded items**. The sampling of the prey in the environment and in the gut was done with two different methodologies (i.e. van Veen grabs and stomach analysis, respectively). To ensure that the sampling of the prey in the environment was representative of the fish diets, we estimated the proportion of prey species biomass found in stomachs that corresponded to species found in the environment. To do so, we associated a benthic community with a fish community only if their relative sampling occurred in the same location (same trawling route) with a temporal distance lower than 31 days (i.e. Gauzens et al., in prep, available as a file for reviewers). Overall, 2487 associations corresponded to this criteria. For a large majority of fish, most of their stomach content matched the results of the environmental sampling (Fig. 4). Overall, for 91% of fish individuals, all of the prey species found in their stomachs were also found in the corresponding environment. For 92% of the fish individuals, more than 90% of the prey biomass observed in their stomachs came from species also observed in the corresponding environment. A limitation of the van Veen grabs is that they might fail species with high mobility. The benthic samples indeed did miss a few fish species that were found in the stomachs of the consumer fish, namely: ‘*Gobiidae spp*.’, ‘*Limanda limanda*’, ‘*Pomatoschistus minutus*’, and ‘*Pleuronectiformes spp*.’. However, in terms of biomass, these species represented less than 0.05 % of the total biomass of prey found in fish consumer stomachs. Another limitation inherent to stomach content analyses are the potential biases associated with differences in digestion times between different species and identification difficulties (Baker et al. 2013). In the absence of species-specific correction factors, it is still possible to apply more subjective correction factors related to tissue types (e.g. soft versus hard-bodied tissues) combined with sensitivity analyses. However, large enough sample size can also average out these biases (Baker et al…).

## Usage Notes

### Data Extraction

To enhance the usability of the database, we have included an R script called “create_csv_files.R.” This script facilitates the extraction of data from the SQL version of the database and generates .csv files.

### Data Utilisation

Due to variations in methodologies and sampling intensities employed for sampling fish and benthic communities, it is important to exercise caution when utilising this dataset in biodiversity studies. For instance, when examining temporal trends of diversity or its response to biotic conditions, a preliminary analysis is necessary to standardize the data (e.g., rarefaction analyses). However, the strong correspondences observed between prey species found in stomach contents and those in the environment (as shown in Figure 4) indicate that the database is well-suited for analyzing predator-prey interactions.

## Acknowledgements

We are profoundly grateful that Wolf E. Arntz collected and provided the valuable data set from his early work in Kiel Bay that we used in this study. We are also thankful to Astrid Jarre who digitized the stomach content data. BG, UB, BR, TB, ACA, LF and MJ gratefully acknowledge the support of iDiv funded by the German Research Foundation (DFG–FZT 118, 202548816). GK acknowledges funding from the German Academic Exchange Service (DAAD, 57070483). MJ acknowledges funding by the European Research Council (ERC) under the European Union’s Horizon 2020 research and innovation programme (grant agreement no. 677232).

## Author Contributions

GK and UJ compiled the initial data. BG, GK and TB formalised the database and created it. BG wrote the first draft of the manuscript with inputs from GK. LF curated the dataset and managed the data deposition. MJ and ACA designed the map. All authors made substantial revisions and comments to the manuscript.

